# The regenerative potential of adult *Nestin+* cerebellar astroglia is limited compared to in neonates

**DOI:** 10.64898/2025.12.01.691510

**Authors:** N. Sumru Bayin, Daniel N. Stephen, Richard Koche, Alexandra L. Joyner

## Abstract

There is a crucial need for strategies that stimulate repair in the adult brain. The neonatal mouse cerebellum can regenerate via the adaptive reprogramming of nestin (Nes)-expressing progenitors (NEPs). However, analysis of *Nes*+ cells of the adult cerebellum is limited. Using reporter lines and genetic inducible fate mapping, we show that adult *Nes*+ cells are mainly Bergmann glia (*Nes*+ Bg) that have *in vitro* sphere-forming ability. Following injury, *Nes*+ Bg increase in number due to upregulation of *Nes* in *Hopx*-expressing Bg, but the cells exhibit limited regeneration. ATAC-seq of *Nes*+ Bg reveals that silencing of developmental genes compared to neonatal NEPs contributes to the impaired regeneration. Activating sonic hedgehog signalling augments the number of *Nes*+ Bg after injury but not neurogenesis, showing additional cues are required. Our results demonstrate an age-dependent decline in the regenerative potential of NEPs and highlight *Nes+* Bg as potential injury-responsive cells that could facilitate regeneration.

## Introduction

The mammalian adult brain has limited regenerative potential upon injury or neuron loss, especially outside of known neurogenic niches. Therefore, it is imperative to study how different cell types in the brain respond to injury and whether they have stem-like characteristics, which in turn can be leveraged to facilitate repair. Previous research has demonstrated that quiescent astroglia take on a stem-like phenotype upon injury and participate in the repair processes^1^. However, the knowledge is limited as to whether other specialised astroglia outside of the known neurogenic niches in the forebrain have stem-like characteristics, and how they react to injury.

In contrast to the limited regeneration in the adult mammalian brain, the neonatal mouse cerebellum is highly regenerative and has emerged as a powerful model to study cellular and molecular mechanisms of regeneration^2–5^. The cerebellum is a key hindbrain structure that is important for motor function, as well as social and cognitive behaviours^6–8^. The cerebellum develops later than the rest of the brain, with its major growth occurring postnatally (2 weeks in mice, up to 6 months in humans)^9,10^, eventually generating the majority of the cells in the brain^11,12^. The cerebellum consists of a central vermis, flanked by two hemispheres and is organised in a folded laminar structure surrounding a core with bilateral nuclei that house the cerebellar output neurons. Given its layered cytoarchitecture and distributed circuitry, regenerative approaches that involve transplantation will be challenging. Therefore, it is crucial to understand how the endogenous mechanisms can be stimulated.

All the cells of the cerebellum are derived from two distinct embryonic progenitor zones. The rhombic lip gives rise to the excitatory neurons, whereas the ventricular zone gives rise to the inhibitory neurons and glia^9,13^. During postnatal development, the rhombic lip-derived granule cell progenitors (GCPs) cover the surface of the cerebellum and continue to proliferate and generate the excitatory granule cells^14–16^. In parallel, several populations of ventricular zone-derived nestin-expressing progenitors (NEPs) located below the GCPs give rise to inhibitory neurons or astrocytes and Bergmann glia (Bg, a specialised astroglia in the cerebellum) via molecularly distinct *Ascl1-*expressing neurogenic or *Hopx-*expressing gliogenic subtypes, respectively^3,4,17,18^. Sonic hedgehog (SHH) secreted from the Purkinje cells is crucial for the proliferation of both progenitor pools and, therefore, for the growth of the cerebellum and scaling of the proportions of several cerebellar cell types^19–23^.

Our previous work showed that upon injuries at birth to Purkinje cells or GCPs, the neonatal cerebellum can replenish the lost cells efficiently^2–4^. The regeneration of GCPs involves adaptive reprogramming of *Hopx-*expressing gliogenic NEPs upon injury^3,4^. When the proliferating GCPs are ablated at birth via irradiation or genetic targeting of diphtheria toxin (DT), the normally gliogenic-NEPs that reside alongside the Purkinje cells in the Bg layer (BgL) first proliferate, then undergo a gliogenic to neurogenic fate switch, and finally migrate to the site of injury and differentiate into GCPs^4^. Analysis of molecular regulators of adaptive reprogramming showed that SHH is required for the proliferation of NEPs during adaptive reprogramming^4^. On the other hand, the injury-induced fate switch of *Hopx-*expressing gliogenic NEPs is regulated by ASCL1, a basic HLH transcription factor that is a known neurogenic factor and used for the direct reprogramming of astrocytes into neurons through ectopic activation^3^. However, whether the adult cerebellum has stem-like cells that express nestin *(Nes)* or *Hopx* and can undergo adaptive reprogramming is unknown.

*Nes* is a well-studied marker of neural stem/progenitor cells during development of many tissues, and also marks the neural stem cells in adult brain neurogenic niches^24,25^. Importantly, *Nes* expression increases in astrocytes and other parenchymal cells, such as ependymal cells around the lateral ventricles upon injury to the cerebral cortex. Increased *Nes* expression is also a hallmark of reactive gliosis^26–29^. These findings highlight that *Nes* expression in other regions of the brain might identify cells with potential stem/progenitor-like properties and/or responsiveness to injury. In the adult cerebellum, Bg continue to reside alongside the Purkinje cells with their cell bodies sitting next to the Purkinje cell soma in the BgL (also known as the Purkinje cell layer) and they extend processes to the pial surface. During postnatal development, Bg processes are required for granule cells to migrate inward to form the internal granule layer (IGL) and they promote the radial migration of Purkinje cells grafted into the adult cerebellum^30,31^. A previous report showed that a rare neurogenic *Nes+* Bg exists in the adult mouse cerebellum^32^. However, to what extent these cells exhibit stemness, their lineage propensity and response to injury remains to be studied.

Here, using transgenic mouse models to identify *Nes+* cells and genetic inducible fate mapping (GIFM), we characterised the distribution of *Nes+* cells throughout the adult mouse cerebellum, across layers and within the anterior-posterior and mediolateral axis, and investigated their lineage relationship to *Hopx*+ cells. We also reported expression of mature astrocyte markers in *Nes*+ cells and their *in vitro* sphere formation ability as a readout for a stem-like characteristic. In addition, we assessed their lineage propensities at homeostasis and upon injury to the adult cerebellum using two different injury models. Importantly, we compared the chromatin accessibility of the adult *Nes*+ cells to their neonatal counterparts, NEPs, and identified potential roadblocks for adaptive reprogramming. Finally, given the importance of SHH during neonatal cerebellum development and regeneration, we tested whether the activation of the pathway upon injury to the adult cerebellum can stimulate *Nes+* cells and facilitate regeneration. In conclusion, our results provide a detailed characterisation of the *Nes*+ cells during homeostasis and upon injury to the adult cerebellum, and offer molecular insights into the limited regenerative potential of the adult brain.

## Results

### *Nes+* cells in the adult cerebellum reside primarily in the BgL

As a first step in studying the *Nes+* cells reported to reside in the adult mouse cerebellum^32^, we performed a detailed characterisation of *Nes+* cells across the medial-lateral and anterior-posterior axes using a *Nes-Cfp* fluorescent reporter line (Figure 1A-D). The transgene (*Nes-Cfp*) expresses cyan fluorescent protein (CFP) under the control of a rat *Nes* promoter^4,33^. Quantification of CFP-expressing cells (Nes-CFP+ cells) on sagittal sections across the medial-lateral axis of 4-6 week old cerebella showed similar densities of Nes-CFP+ cells in the different layers based on medial-lateral location (Figure 1E, n=3 brains) and that the majority of the Nes-CFP+ cells were in the BgL (82.44 ± 5.93, n=3 brains, Figure 1F). Across the anterior-posterior axis, the majority of the Nes-CFP+ cells were also localised in the BgL (Figure 1G). Furthermore, there was a significantly higher proportion of Nes-CFP+ cells in the central (lobules 5-8) and posterior (lobules 9, 10) sectors compared to the anterior (lobules 1-5) sector, with the highest proportion in the posterior sector when normalized to section area (Figure 1G). Finally, to further study the identity of these cells, we assessed whether Nes-CFP+ cells co-expressed SOX2 and/or S100β. SOX2 has been reported to mark neural progenitors and NEPs in the early postnatal cerebellum and some adult cerebellar astroglia, whereas S100β is restricted to mature Bg and astrocytes at all stages. Analysis of SOX2 and S100β co-expression with Nes-CFP revealed that the majority of Nes-CFP+ cells were positive for SOX2 (97.15 ± 1.48%, n=3) and very few of these cells expressed S100B (14.27 ± 12.25%, n=3), suggesting an immature/progenitor-like phenotype of Nes-CFP+ cells (Figure 1H). Furthermore, only rare Nes-CFP+ cells were negative for both markers (2.85 ± 1.48%, n=3, Figure 1H). Interestingly, the SOX2+ S100β-Nes-CFP+ cells showed a unique localisation between the two main layers of Bg in the medial-lateral axis, highlighting their possible unique identity compared to mature Bg (Figure 1I).

**Figure 1.**
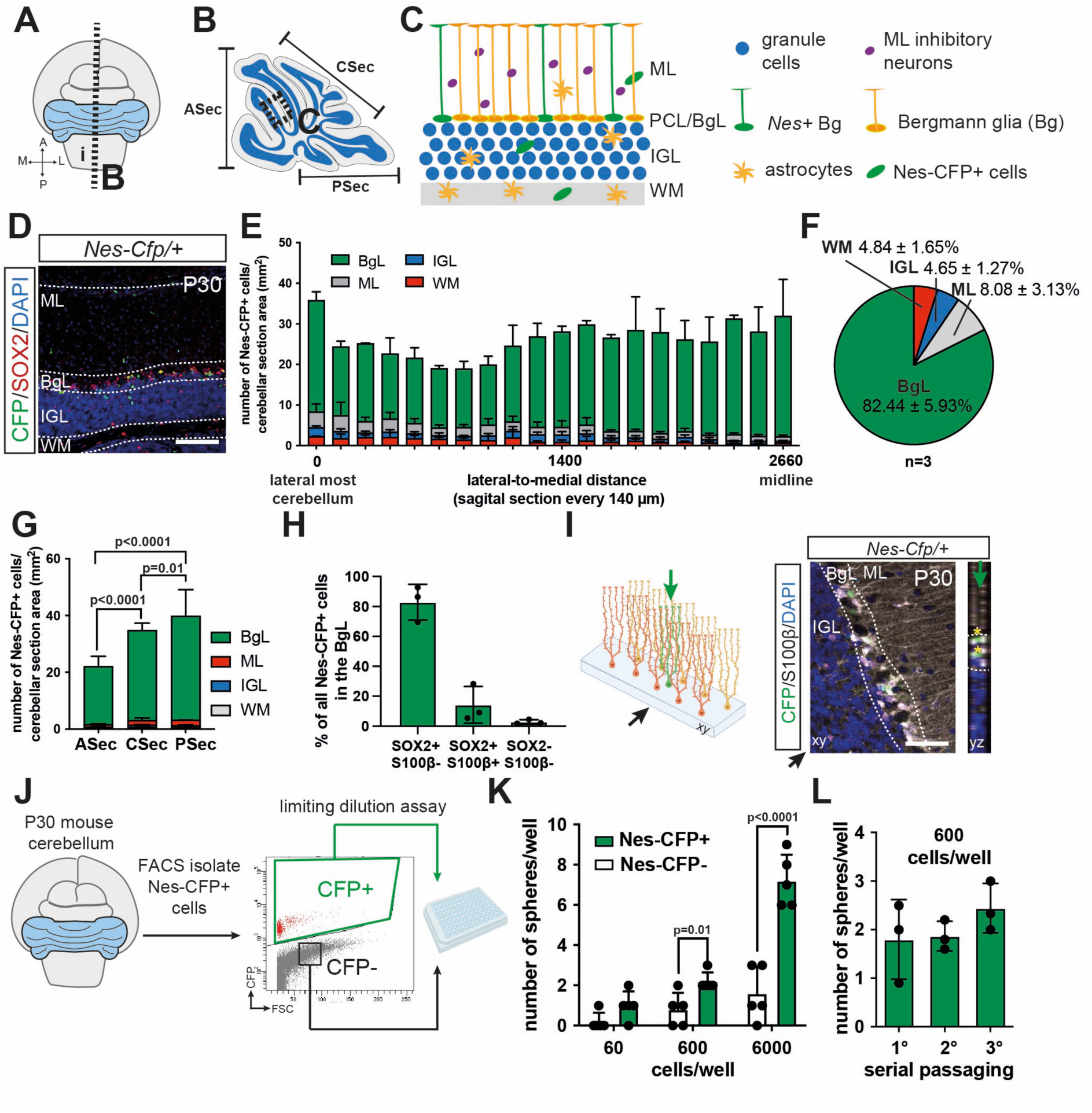
*Nes*+ cells in the adult cerebellum reside primarily in the BgL. **A-C)** Schematics of the adult cerebellum and cytoarchitecture. **D)** Immunofluorescent analysis of P30 *Nes-Cfp/+* animals showing the localisation of Nes-CFP+ cells across the layers of the cortex. **E-F)** Quantification of the density of Nes-CFP+ cells in different layers of the cerebellum across the medial-lateral axis (E), on average, 82% of the cells reside in the BgL (F, n=3). **G)** Quantification of the density of Nes-CFP+ cells across the anteroposterior axis on midsagittal sections shows a significantly higher density in the central and posterior sectors than anterior sector, with the posterior sector having the greatest density of Nes-CFP+ cells in the BgL (Two-way ANOVA, F_(2,16)_=18.6, p<0.0001, n=3). **H)** Analysis of the percentage of SOX2+ and/or S100β+ Nes-CFP+ cells. **I)** Schematic (left) and orthogonal projection of a confocal microscopy z-stack showing the unique localisation of Nes-CFP+ cells within the BgL. Black arrow shows the xy-axis and the green arrow shows the yz-axis in the schematic and the image. **J-L)** FACS-isolated Nes-CFP+ cells form neurospheres more efficiently than the Nes-CFP-cells (Two-way ANOVA, F_(1,4)_=39.5, p=0.003, n=5), and their sphere-forming capacity is maintained across serial passaging (at 600 cells/well density). Multiple comparison tests are shown in the figures. Scale bars: D: 100 µm, I: 50 µm.

### *Nes+* adult cerebellar cells have stem/progenitor cell-like characteristics *in vitro*

To assess whether *Nes+* cells of the adult cerebellum have stem/progenitor cell potential, we performed a limiting dilution neurosphere formation assay on freshly isolated Nes-CFP+ cells from the cerebellum of 4-6 weeks-old mice. Fluorescence-assisted cell sorting (FACS) was used to isolate CFP+ and CFP-cells, and the cells were plated at limiting dilutions (60, 600 and 6000 cells/well) to form spheres under self-renewing conditions with EGF and bFGF (Figure 1J). Nes-CFP+ cells formed significantly more spheres compared to the CFP-cells when plated at 600 and 6000 cells/well densities (Figure 1K). Furthermore, the neurosphere-forming ability of the Nes-CFP+ cells was maintained over serial passaging (Figure 1L), suggesting long-term self-renewal potential, albeit of rare Nes-CFP+ cells. Collectively, the data so far suggest that the majority of the Nes-CFP+ cells in the cerebellum represent a unique population of Bg cells based on their location and stem/progenitor cell-like property *in vitro*, and lack of the mature astrocyte marker S100β.

### Nes-CFP+ Bg numbers are increased in response to injury

Our previous research highlighted that the neonatal cerebellum is highly regenerative and can recover efficiently from the loss of two cell types at birth^2–4^. Given that the adult Nes-CFP*+* Bg possess some stem/progenitor cell-like features, we hypothesised that they can facilitate regeneration and repair upon injury to the adult cerebellum. To test this, we developed two injury paradigms. In one paradigm, we used a photothrombotic local stroke model to induce a vermis-targeted stroke (ischemia) (Figure 2A)^34^. In the other paradigm, we utilised a pharmacogenetic approach, wherein the granule cells are selectively depleted in a transgenic mouse model in which the granule cells express a diphtheria toxin receptor (DTR) and are exposed to DT. *Atoh1-Tta; TRE-Cre; R26^lox-STOP-loxDTR^* animals were given doxycycline (DOX) between embryonic day (E) 8.5 and 12.5 to prevent TTA activity and Cre-mediated recombination to induce DTR expression in the excitatory cerebellar nuclei neurons that transiently express *Atoh1* at this stage (referred to as *GC-DTR* mice)^4^. The cell ablation was then performed via systemic injection of DT in 4-5 week-old mice (Supplementary Figure 1A-B). While the ischemia model provides a physiological injury, *GC-DTR* allows for comparative experiments between neonatal and adult injury to the granule cell lineage. Both injuries were performed on *Nes-Cfp* mice to detect Nes-CFP*+* Bg and assess their immediate response to injury.

**Figure 2.**
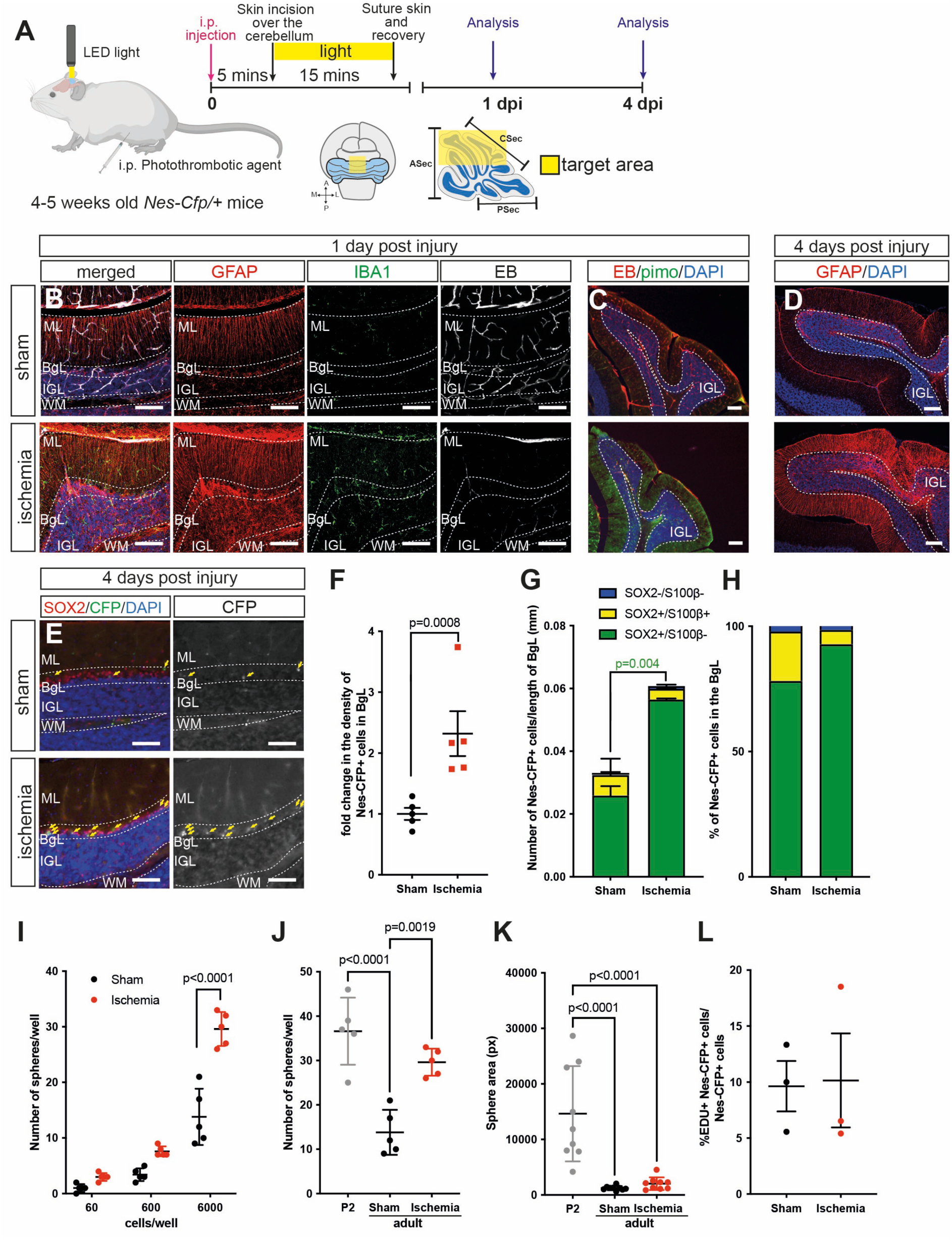
Nes-CFP+ Bg density is increased in response to ischemic injury to the adult cerebellum. **A)** Schematic explaining the injury paradigm. **B-D)** Analysis of sections from sham or ischemic cerebella 1 day post injury shows increased microglia density (IBA1+ cells), reduced vascular perfusion assessed by intravenous Evans blue (EB) injection and increased hypoxia in the injured regions based on pimonidazole (pimo) (C). Gliosis can be observed shortly after injury with increased GFAP levels, 4 days post injury (D). **E-H)** Analysis of Nes-CFP+ cells in the central sector (lobules 6-8) shows a significant increase in the density of Nes-CFP+ cells in the BgL after injury, most of which are SOX2+ and S100β-(Student’s t-test, F: p<0.0008, G: p<0.004, n=5). **I-K)** FACS isolated Nes-CFP+ cells from sham or ischemic cerebellar (vermis only) show a significant increase in the in vitro neurosphere forming ability at a density of 6,000 cells/well (I: Two-way ANOVA, F_(1,4)_=33.1, p=0.0045, n=5), which is similar to neonatal Nes-CFP+ cells (6,000 cells/well) isolated from P2 cerebella (J: One-way ANOVA, F_(2,12)_=22.17, p<0.0001, n=5). However, the size of the neurospheres generated from either adult sham or ischemic cerebellar was significantly smaller than neonatal spheres (One-way ANOVA, F_(2,24)_=20.26, p<0.0001, n=9). **L)** Analysis of EdU+ cells, when EdU was injected 1 hour before sacrifice 4 days post adult injury does not show a significant change in the % EdU+ Nes-CFP+ cells upon injury (Student’s t-test, p>0.05, n=3). Multiple comparison tests are shown in the figures. Scale bars: 100 µm

Using the ischemic model, we first investigated the changes in the brain microenvironment to confirm a stroke-like phenotype. Immunofluorescent analysis of the cerebella from control (Sham) and injured (Ischemia) mice one day after injury showed a dramatic reduction in the vascular perfusion of blood, assessed via intravenous Evans Blue injection (Figure 2B). Pimonidazole staining showed an increase, which indicates increased hypoxia in the targeted area (Figure 2C). In addition, we observed increased microglia throughout the injured cerebellar lobules along with an increase in GFAP intensity 1 day after injury (Figure 2B). The increase in GFAP levels, especially in the molecular layer and IGL, was sustained at 4 days after injury, demonstrating reactive gliosis at the site of injury (Figure 2D). Thus, an injury response was mounted after ischemia to the cerebellum.

Next, we tested whether a response to injury involves a change in the density of Nes-CFP+ Bg by quantifying their numbers. Analysis of Nes-CFP+ Bg density 4 days after injury at the site of injury (vermis, central sector) revealed a 2.32 ± 0.82-fold (n=5) increase in cell density (Figure 2F). Importantly, the majority of the increase was driven by SOX2+ S100β-cells, suggesting an increase in the progenitor-like cells (Figure 2G-H). The density of the Nes-CFP+ cells in the molecular layer, IGL and WM were highly variable and not significantly changed following injury (Supplementary Figure 1F). Importantly, the lateral cerebellum away from the site of injury did not show an increase in the density of Nes-CFP+ cells in the BgL, further highlighting that the increase in the Nes-CFP+ Bg is specific to the ischemia (Supplementary Figure 1G).

Similarly, when the granule cells were ablated in *GC-DTR* animals via DT injection at P30, we observed an increase in microglia density in the IGL, indicating an injury-like phenotype 1 day after DT injection, and the density of Nes-CFP+ Bg showed a 4.59 ± 2.63-fold increase (n=4) 4 days after the ablation (Supplementary Figure 1C-D). Finally, the increase in the number of Nes-CFP+ Bg was again driven by an increase in the SOX2+ S100β - cells (Supplementary Figure 1E), suggesting an increase in the stem/progenitor-like cells upon injury. In summary, Nes-CFP+ Bg are the main *Nes*-expressing cell type in the adult cerebellum that responds to proximal cell loss by increasing their numbers.

### Nes-CFP+ cells form more neurospheres post injury

Previous reports have shown that injury can stimulate astrocytes to become more stem cell-like^35–37^. To test whether injury changes the *in vitro* self-renewal potential of the *Nes+* Bg, we FACS-isolated Nes-CFP+ cells from the vermis of sham or ischemic cerebella 4 days after injury and subjected them to a limiting dilution neurosphere formation assay. Nes-CFP+ cells from ischemic animals formed approximately 2-fold more neurospheres than counterparts isolated from sham animals at 6000cell/well density (n=5, Figure 2I). Interestingly, ischemia increased the neurosphere formation capacity to levels similar to the level of neonatal Nes-CFP+ cells isolated from P2 *Nes-Cfp* cerebella (n=5) (Figure 2J). However, when the area of the neurospheres was quantified, which indirectly represents the proliferation capacity of the cells, we observed that the spheres generated from adult sham and ischemic Nes-CFP+ cells were significantly smaller than the ones generated from neonatal Nes-CFP+ (Figure 2K). In line with this observation, when we analysed the proliferation of Nes-CFP+ Bg via EdU injection 1 hour before analysis of adult cerebella from sham and ischemic mice 4 days after injury, although the total number of EdU+ Nes-CFP+ Bg increased, we observed that the percentages of Nes-CFP+ Bg that were EdU+ remained unchanged (Figure 2L), suggesting no increase in their proliferative potential 4 days after injury. In summary, these results suggest that although injury leads to an increase in the density of Nes-CFP+ Bg and their *in vitro* neurosphere formation ability, their proliferation ability remains limited.

### Comparisons of the chromatin landscape between the neonatal and adult Nes-CFP+ cells suggest silencing of pro-regenerative mechanisms

Guided by our knowledge of the cellular behaviours during neonatal cerebellar regeneration, an increase in the proliferation of the regenerative cells is needed to replenish the lost cells upon injury. For example, during regeneration of GCPs upon injury to the cerebellum at birth, shortly after GCP cell death, *Hopx*-expressing gliogenic NEPs in the BgL significantly increase their proliferation, prior to undergoing an *Ascl1*-mediated fate switch and migration to the EGL^3^. In contrast, upon injury to the adult cerebellum, although we observed an increase in the density of Nes-CFP+ Bg, their proportion of cells undergoing mitosis was limited and not enhanced with injury (Figure 2L). We hypothesised that this could be due to the limited availability of pro-proliferative signals in the adult cerebellum compared to the neonate and/or the silencing of key stemness, proliferation and neurogenic genes once the developmental window ends. To test this hypothesis, we investigated chromatin accessibility via ATAC-seq on FACS-isolated neonatal and adult Nes-CFP+ cerebellar cells 4 days after injury from control and injured *GC-DTR* cerebella that were injected with DT at P1 or P30, respectively (Figure 3A). The *GC-DTR* model was chosen for this experiment to ensure the analysis was performed on the same injury paradigm and genetic background at the two ages.

**Figure 3.**
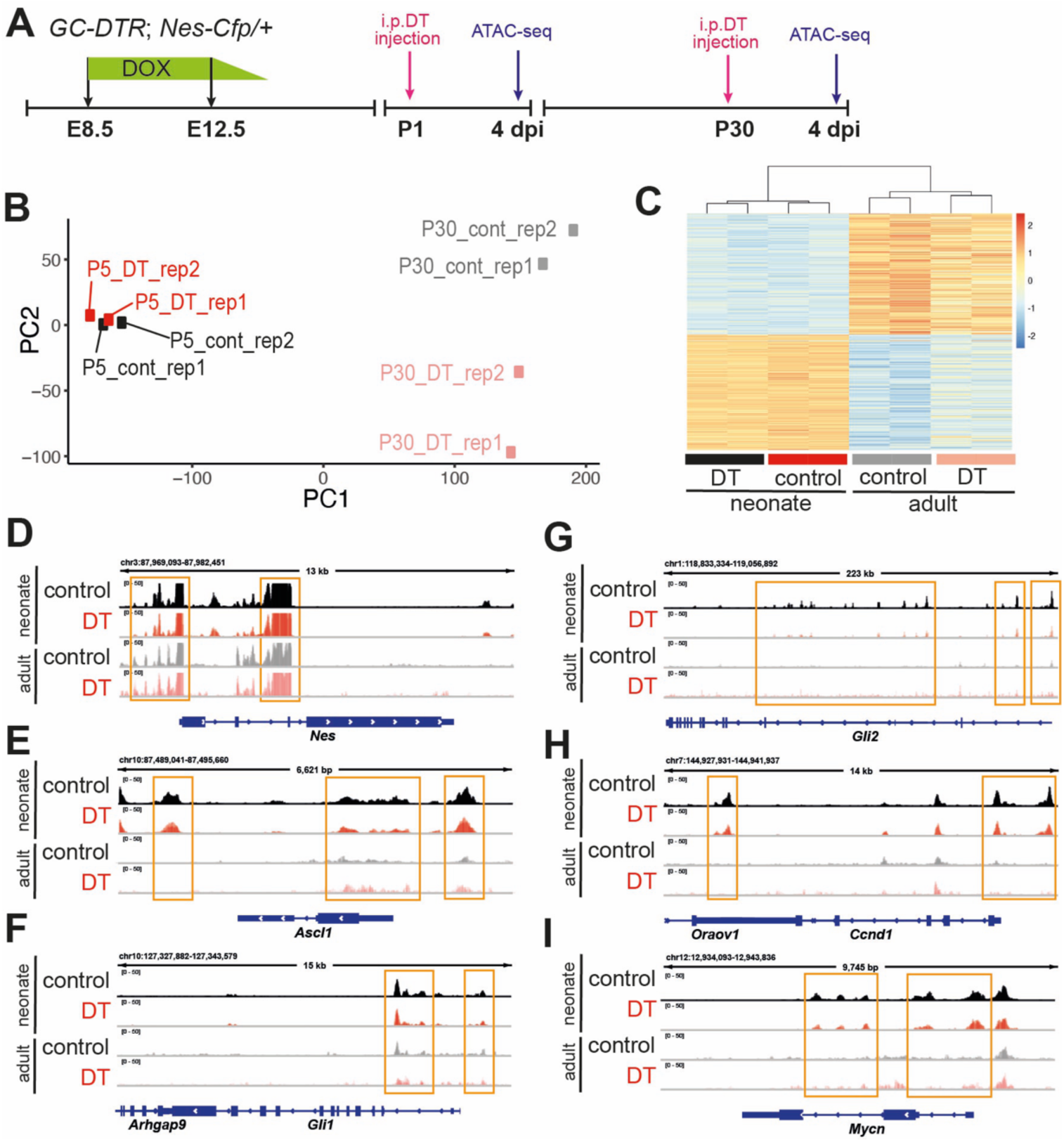
The chromatin landscape is dramatically different between P5 and P30 Nes-CFP+ cells. **A)** Experimental setup for bulk ATAC-seq analysis of FACS-sorted Nes-CFP+ cells from neonatal and adult, control or injured *GC-DTR* mice that were given DT at P5 or P30 and analysed 4 days after injection. **B)** Principal component analysis (PCA) of the samples shows that the main differences are driven by age. **C)** Hierarchical clustering of all significant differentially open peaks identified from the comparison of P5 to P30 control Nes-CFP+ cells was performed on all conditions: the control and injured Nes-CFP+ cells at P5 or P30 (see Supplementary Table 2). **D-I)** IGV tracks of genes of interest that are involved in SHH signalling pathway (*Gli1/2, Ccnd1, Mycn*) or adaptive reprogramming (*Ascl1*) that show changes in chromatin accessibility in the adult Nes-CFP+ cells compared to the neonates (p_adj<0.1, Supplementary Table 2). Accessibility around *Nes* is observed in all conditions.

Principal component analysis (PCA) revealed that the predominant changes in the chromatin accessibility were driven by age (PC1) and then upon injury to the adult cerebellum (PC2) (Figure 3B). However, we did not observe significant differences upon injury in the chromatin accessibility of Nes-CFP+ cells at P5 (4 days after DT injection) compared to controls via bulk ATAC-seq. This result contrasts with our previous analysis at P2 of cerebella irradiated at P1, where we observed increased accessibility of stress response genes shortly after injury to the EGL^5^. Such bulk analyses may be confounded by contrasting changes in different NEP subpopulations, which are difficult to distinguish via a bulk approach and also the proportion of NEPs in the BgL at P5 is likely lower than at P2. On the other hand, the adult Nes-CFP+ cells showed significant changes in chromatin accessibility 4 days after injury at P30 compared to their controls (Supplementary Figure 2 and Supplementary Table 1). Gene ontology (GO) analysis of the genes associated with differentially open chromatin upon injury to the adult cerebellum revealed GO-terms related to organ growth and nervous system development (n=575 genes, Supplementary figure 2A), whereas the genes associated with open chromatin in control adult Nes-CFP+ cells were associated with various cellular processes involved in calcium signalling, synaptic processes or cytoskeleton organisation (n=160 genes, Supplementary Figure 2B). These results are supportive of our observation that adult Nes-CFP+ Bg take on a more “stem-like” phenotype upon injury, as exhibited by the increased *in vitro* sphere-forming ability (Figure 2J).

As suggested by the PCA, the predominant changes in the chromatin accessibility are driven by age, likely highlighting the epigenetic changes that occur once development is over. To provide support for this idea, we identified the significant differentially accessible chromatin between the P5 and P30 Nes-CFP+ cells from the control brains and compared these regions to the accessible chromatin from neonatal and adult, control and injured Nes-CFP+ cells (Figure 3C, Supplementary Table 2). Hierarchical clustering of these regions in control and injured Nes-CFP+ cells from both neonatal and adult cerebellum also showed that the samples clustered by age (Figure 3C), further confirming the age-driven changes in the chromatin landscape suggested by the PCA analysis (Figure 3B).

We next interrogated the data in a locus-specific manner. Firstly, the open chromatin identified around the *Nes* promoter confirmed the NEP identity of Nes-CFP+ cells isolated (Figure 3D). Given that neonatal NEP proliferation is stimulated by SHH signalling and proliferation of NEPs during regeneration is dependent on SHH signalling^4^, we assessed chromatin accessibility in our bulk ATAC-seq around some of the SHH pathways mediators and target genes (*Gli1, Gli2, Mycn, Ccnd1, cMyc*) that are involved in cellular proliferation and also the proneural transcription factor *Ascl1* due to its requirement for adaptive reprogramming^3^ (Figure 3E-I). We found reduced chromatin accessibility around these genes in the adult Nes-CFP+ cells from control cerebella compared to the neonatal cells from controls (Figure 3E-I, Supplementary Table 2). The chromatin accessibility around these genes did not differ between the control and injured cerebellum at both ages (Figure 3E-I). In summary, the board chromatin accessibility in the neonatal Nes-CFP+ cells, with and without injury, compared to the adult Nes-CFP+ cells around some genes involved in proliferation and the SHH pathway suggests permissiveness of the neonatal NEPs to regeneration and adaptive reprogramming in contrast to the adult Nes-CFP+ Bg. These results also suggest that reduced SHH pathway activity in Nes-CFP+ cells could be one of the reasons behind the limited injury response of *Nes+* Bg upon injury to the adult cerebellum.

### Activation of SHH signalling augments the injury response of *Nes+* Bg by increasing the number of Nes-CFP+ Bg

Given the importance of SHH during neonatal NEP adaptive reprogramming^4^ and the reduced chromatin accessibility around its target genes in the adult Nes-CFP+ cells, we reasoned that activating the SHH pathway could stimulate *Nes+* Bg and therefore enhance their regenerative potential. To test this idea, we analysed sections from *Shh-P1* transgenic mouse cerebella, which bear a 3^rd^ copy of the *Shh* gene and thus have increased SHH signalling in the cerebellum^22^. Analysis of Nes-CFP+ cells in the vermis of *Shh-P1* mice showed a significant increase in the density of Nes-CFP+ cells in the BgL (Supplementary Figure 3A-C), supporting that *Nes+* Bg cells are responsive to SHH signalling.

We then tested whether a small molecule agonist of SMO (SAG) that activates SHH signalling increases the density of Nes-CFP+ cells. We first assessed whether 10 µg/g systemic injection of SAG activates SHH signalling using a *Gli1^eGFP/+^*reporter line^38^ and quantified *Gli1* expression via qRT-PCR on whole cerebellum extracts and GFP intensity on cerebellar sections (Supplementary Figure 3D-F). Six hours after injection, we observed a significant increase in both the *Gli1* expression (1.82 ± 0.29-fold, n=3 brains) and eGFP intensity (1.91 ± 0.16-fold, n=3) in SAG treated cerebella compared to the vehicle injected animals, suggesting that this concentration of SAG is active in the brain after systemic injection, consistent with previous work^39^.

Next, we tested whether SAG treatment *in vivo* improves the *in vitro* self-renewal capacity of Nes-CFP+ Bg. 4-5 week old *Nes-Cfp*+ animals were injected with three 10ug/g SAG injections every 24 hours and Nes-CFP+ cells were isolated by FACS. The limiting dilution neurosphere formation assay was then performed (Figure 4A). We observed a significant increase in the number of neurospheres formed with Nes-CFP+ cells isolated from SAG-injected animals when plated at high density (6,000 cells/well, Figure 4B), suggesting an increase in the *in vitro* self-renewal capacity. This was previously observed in cortical astrocytes isolated from injured brains that were administered SHH agonist^37^.

**Figure 4.**
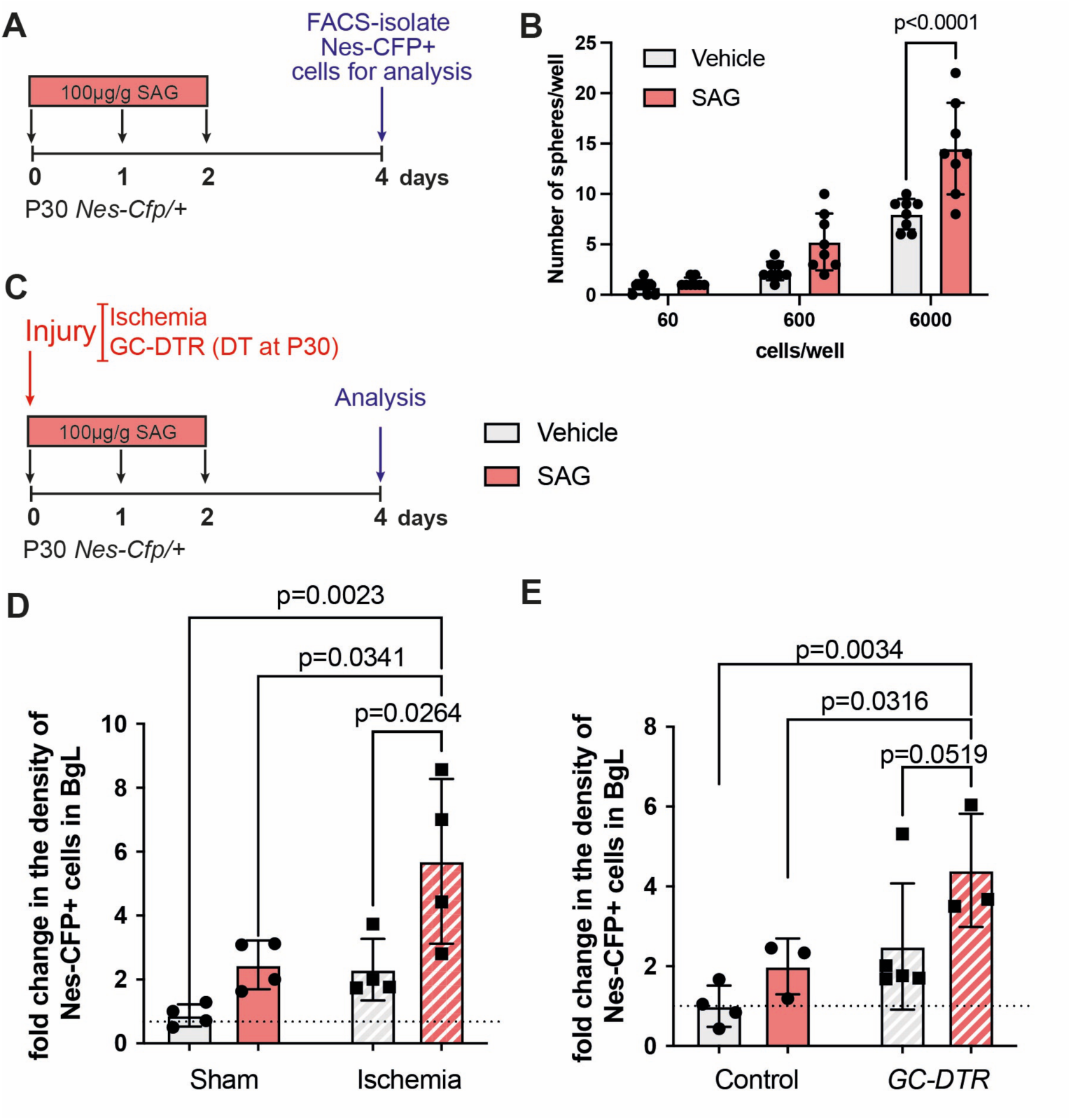
Activation of SHH signalling augments the injury response of *Nes+* Bg by increasing the number of Nes-CFP+ Bg and neurosphere formation. **A)** Schematics of experimental timeline. **B)** FACS isolated Nes-CFP+ cells from vehicle or SAG injected animals show SAG increases neurosphere forming ability, in particular at 6,000 cells/well density (Two-way ANOVA, F_(1,42)_=24.12, p<0.0001, n=8). **C)** Schematics of experimental timeline. **D-E)** Analysis of fold-change in the Nes-CFP+ cell densities in the BgL in ischemic (D: SAG treatment: Two-way ANOVA, F_(1,12)_=11.97, p=0.0047, n=4; Injury: Two-way ANOVA, F_(1,12)_=10.59, p=0.0069, n=4) and pharmacogenetic injury model (E: SAG treatment: Two-way ANOVA, F_(1,11)_=5.248, p=0.042, n≥3; Injury: Two-way ANOVA, F_(1,11)_=9.506, p=0.01, n≥3). Multiple comparison tests are shown in the figures.

Next, we activated the SHH pathway after injury to the adult cerebellum in both the ischemic injury and *GC-DTR* models and quantified the density of Nes-CFP+ Bg with and without SAG administration (Figure 4C). We observed a significant, approximately two-fold increase in the density of Nes-CFP+ Bg upon SAG administration, similar to the increase with ischemic injury alone compared to sham animals with vehicle treatment (Figure 4D). Importantly, the increase in the density of Nes-CFP+ Bg 4 days after injury was augmented further when SAG was combined with ischemia, suggesting an additive effect (Figure 4D). A similar increase in the density of Nes-CFP+ Bg was also observed in *GC-DTR* animals that were administered SAG compared to no SAG (Figure 4E). Finally, no significant changes were observed in the other Nes-CFP+ cells and the proportions of SOX2+ astroglia in the other layers of the cerebellum in SAG compared to vehicle-treated sham or ischemia animals (Supplementary Figure 3G-I).

Finally, to test whether the increase in the Nes-CFP+ Bg densities following SAG was due to increased proliferation, we administered EdU daily starting one day after injury (Supplementary Figure 3J). Interestingly, although not significant, SAG administration alone or injury alone showed a trend towards an increase in the density of the EdU+ Nes-CFP+ cells in the BgL (Supplementary Figure 3K). However, this effect was not enhanced in SAG in combination with injury, suggesting the involvement of additional mechanisms that increase the density of Nes-CFP+ Bg. Importantly, given the increase in the density of Nes-CFP*+* Bg with SAG and/or injury, the percentages of the Nes-CFP*+* Bg that were proliferating (EdU+) did not change in all conditions (Supplementary Figure 3L). This data indicates that the additive effect of SAG upon injury on the density of Nes-CFP+ cells is not due to an increase in proliferation upon SHH activation. These results also suggest potentially different roles for SHH in adult cerebellum astroglia compared to its mitogenic effects in the neonatal cerebellum and points to a quiescent Bg in the adult that upregulates *Nes* expression following injury, with no effect on proliferation of all *Nes+* Bg.

### *Nes+* Bg generate astroglia during homeostasis and upon injury, but not neurons

A previous study of *Nes+* cells in the BgL suggested that exercise can increase their numbers and that they rarely give rise to new granule cells based on a label retention assay^32^. An important question that remains is whether *Nes+* Bg have any neurogenic potential during homeostasis and, importantly, upon injury. In addition, whether SAG administration can change the lineage propensities of *Nes+* Bg. We therefore performed GIFM using *Nes-FlpoERT2/+; R26^Frt-STOP-FrtTdTom/+^* (*Nes-Td*) mice. Given the low frequency of Nes-CFP+ Bg and to ensure that our results are not confounded by background recombination in *Nes-Td* mice without Tamoxifen, we first analysed *Nes-Td* brains that were not injected with Tamoxifen or were injected with Tamoxifen at P30 and analysed 2, 6 days or 2 weeks after Tamoxifen injection (Supplementary Figure 4A). We observed a significant level of recombination in the absence of Tamoxifen, specifically in molecular layer inhibitory neurons and to a lesser extent in oligodendrocytes and OPC-like cells based on morphology. Importantly, the density of TdTom+ molecular layer inhibitory was similar between no Tamoxifen, and Tamoxifen-injected sham and injured brains (Supplementary Figure 4B-D). Thus, recombination in these cell types must occur during development (before P30). Tamoxifen resulted in labelling of Bg and astrocytes, as well as some perivascular cells that likely represent *Nes+* pericytes. We omitted molecular layer inhibitory neurons, pericytes and oligodendrocyte lineage cells from our downstream analysis and only focused on astroglia due to their relevance and labelling following Tamoxifen treatment. Finally, temporal analysis during homeostasis over the 2 weeks following Tamoxifen administration showed no significant changes in the numbers of TdTom+ astrocytes or Bg, suggesting that at steady state, there is minimal gliogenesis from the *Nes*+ cells in the adult cerebellum (Supplementary figure 4C).

Next, we performed GIFM using sham or ischemic 4-5 week old *Nes-Td* animals that were given 200 µg/g Tamoxifen 4 days after ischemic injury. For mice given SAG, it was administered daily for 3 days as above, starting on the day of surgery (Figure 5A). Analysis of brains from *Nes-Td; Nes-Cfp* mice 2 days after Tamoxifen injection confirmed that TdTom+ Bg were also Nes-CFP+ (Figure 5B), showing that the *Nes* promoters in both transgenes (*Nes-Cfp* and *Nes-FlpoERT2*) label similar populations. In line with this, we observed similar changes to the density of TdTom+ cells in the BgL two weeks after injury, to the density of Nes-CFP+ Bg observed 4 days after injury (Figure 5C compared to Fig. 4D). Consistent with our analysis of Nes-CFP+ Bg density, SAG or injury alone similarly increased the density of TdTom labelled cells in the BgL of *Nes-Td* mice. The combination of both conditions, although did not lead to a significantly greater increase in the density of TdTom+ Bg than either condition alone, showed a trend towards it (Figure 5C). As expected, the density of TdTom+ astrocytes outside the BgL showed no significant changes after injury and/or SAG treatment, and there was no increase in neurons after injury (Figure 5D, Supplementary Figure 4D). In summary, these data suggest that *Nes+* Bg continue to be gliogenic after injury, similar to their neonatal counterparts during homoestasis and are unable to generate neurons upon injury. In addition, these results provide further evidence that *Nes*+ Bg and cerebellar astrocytes have differential responses to SHH activation and injury, and SAG treatment does not promote *Nes+* Bg to generate neurons upon injury.

**Figure 5.**
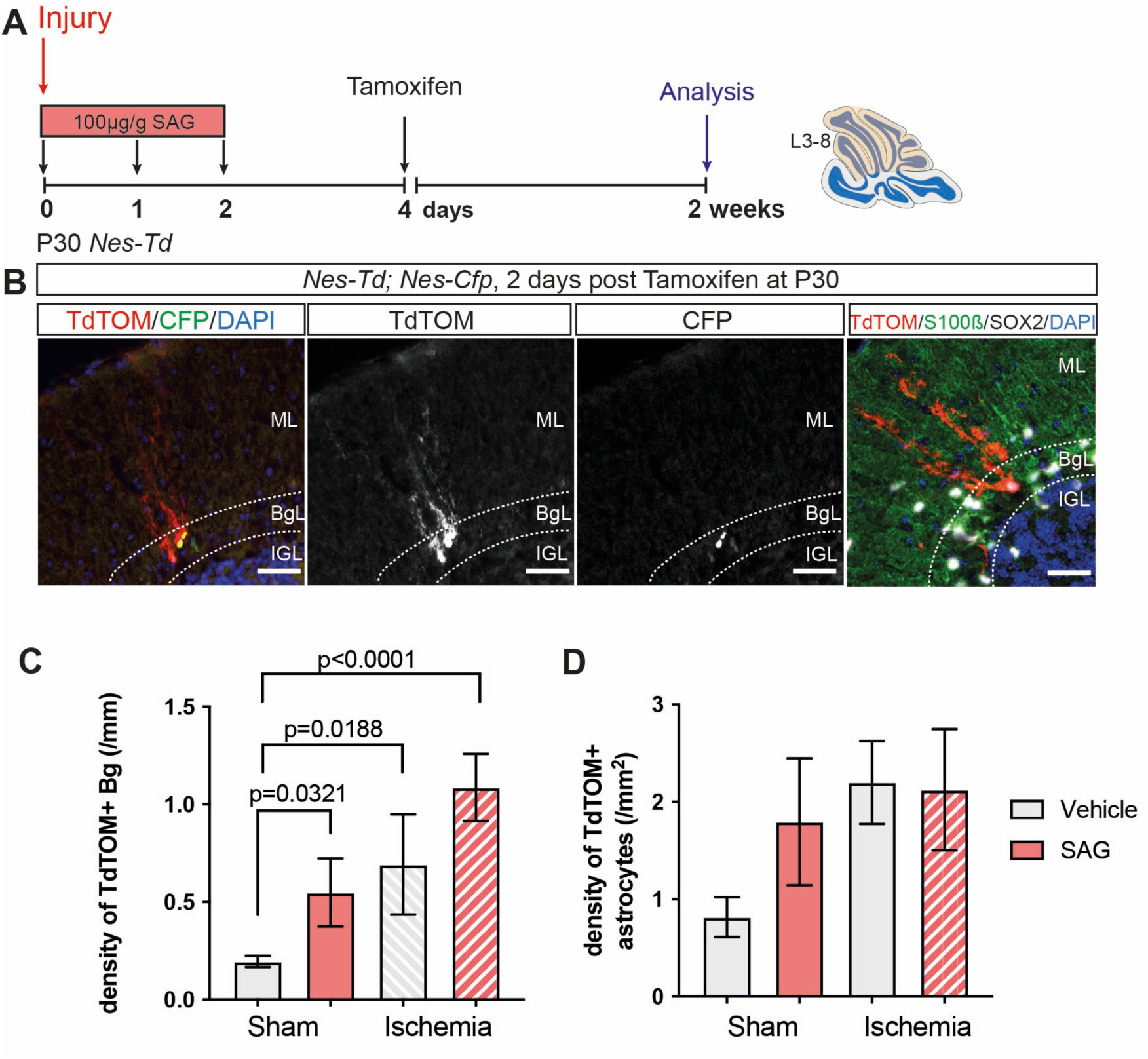
*Nes*+ Bg generate astroglia during homeostasis and upon injury, but not neurons. **A)** Experimental timeline for the GIFM of *Nes-FlpoERT2/+; R26^Frt-STOP-FrtTdTom/+^* (*Nes-Td*) mice. **B)** Labelling of TdTom and CFP in *Nes-Td*; *Nes-Cfp* mice shows double labelled Bg 2 days after Tamoxifen injection. **C-D)** Quantification of the density of TdTom+ Bg (C: One-way ANOVA, F_(3,27)_=6.236, p=0.0023, n≥6, individual Student’s t-test are shown in the figure) and astrocytes outside the BgL (D: One-way ANOVA, F_(3,25)_=1.913, p=0.1532, n≥6) upon injury. Multiple comparison tests are shown in the figures. Data is shown as mean ± SEM. Scale bars: 50 µm

### Injury stimulates *Hopx*+ quiescent Bg to become *Nes*+

While the significant increase in the density of Nes*-*CFP*+* cells in the BgL 4 days after injury and/or SAG, and a similar increase in TdTom+ cells in *Nes-Td* mice (given Tamoxifen at 4 days after injury) show that the density of these cells can increase with each stimulus, the limited proliferation of the cells observed in all conditions suggests that a distinct population of Bg are induced to express *Nes* (and therefore, Nes-CFP) after injury and/or SAG treatment. Consistent with this hypothesis, when we performed GIFM and injected *Nes-Td* animals with Tamoxifen 2 days before ischemia and analysed the brains 2 weeks after injury (Figure 6A), we observed no significant difference in the number of TdTom+ Bg produced between sham and ischemia conditions (Figure 6B). Labelling of astrocytes outside the BgL also did not change in either condition (Figure 6C). These data suggest that the injury induces a group of *Nes-* Bg to become *Nes+*. One possibility is that there is a quiescent SOX2+ Bg population that expresses *Hopx,* a marker of some neonatal gliogenic NEPs and adult Bg^3,40^. To test this hypothesis, we performed GIFM using *Hopx^CreERT^*^2^*^/+^; R26^lox-STOP-loxTdTom/+^*(*Hopx-Td*) animals that were given Tamoxifen 3 consecutive times, 2 days apart, prior to ischemia, to maximise labelling Bg (Figure 6D). We previously reported that *Hopx^CreERT2^* does not result in background recombination in the cerebellum without Tamoxifen injection^3^. Analysis of TdTom+ cells in *Hopx-Td* animals treated with Tamoxifen before injury showed significant labelling in the BgL, along with some astrocytes in the IGL and WM (Figure 6E). Interestingly, 4 days after injury, the ischemic brains showed a significant 1.83 ± 0.60-fold (n=4) increase in the number of TdTom+ cells in the BgL compared to the sham controls (Figure 6F). No significant change in the density of other astrocytes was detected (Figure 6G). Importantly, analysis of the Hopx-Td+ cells in the BgL showed that the majority of TdTom labelled cells were SOX2+ S100β-and the increase after injury was driven by SOX2+ S100β-cells, highlighting their immature phenotype similar to Nes-CFP+ Bg (Figure 6H). Collectively, these results suggest that injury stimulates *Hopx+* Bg to become new *Nes+* Bg (Figure 6I).

**Figure 6.**
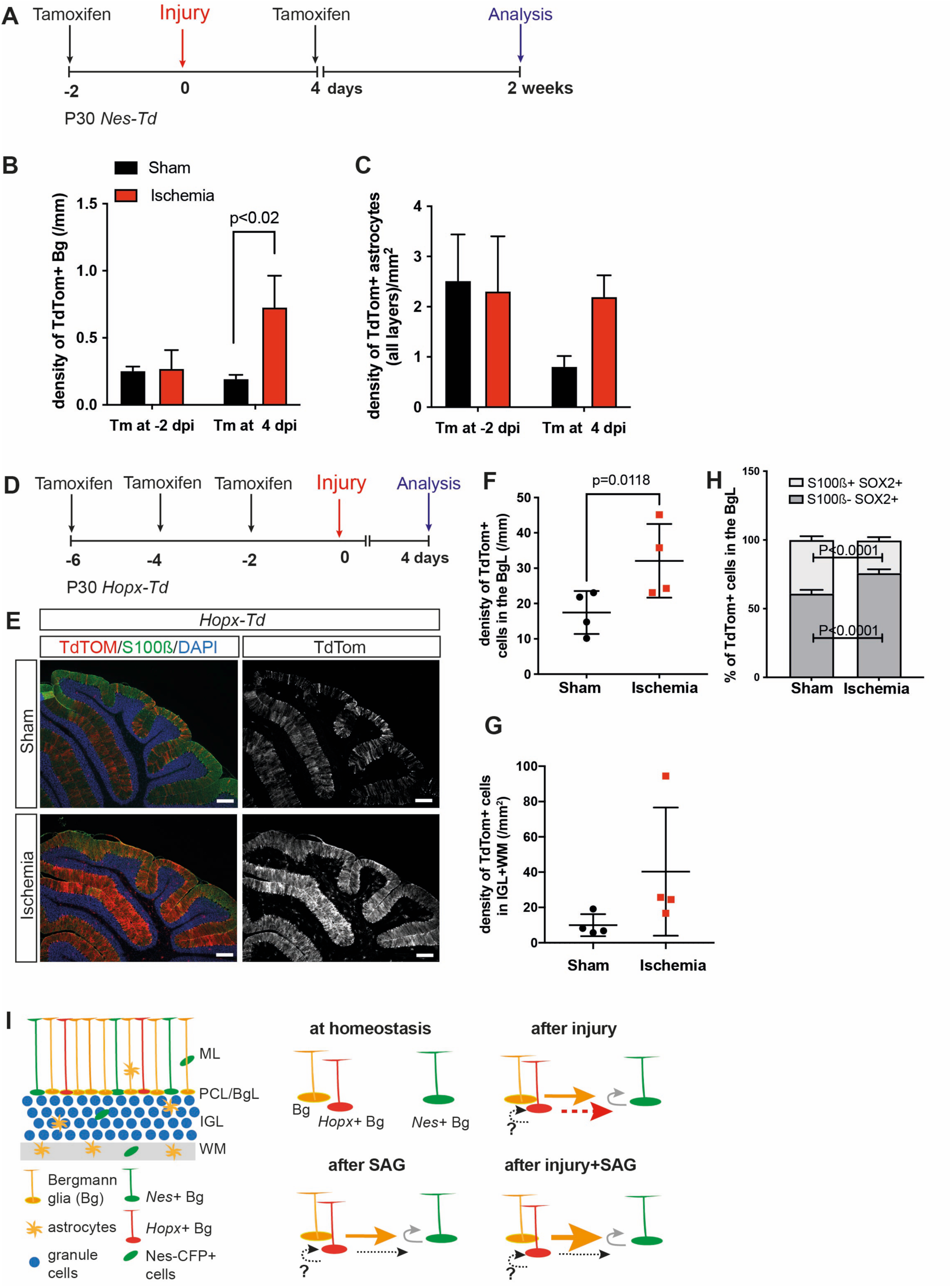
Injury stimulates *Hopx*+ quiescent Bg to become *Nes*+. **A)** Experimental timeline. **B-C)** Quantification of the density of TdTom+ Bg in the BgL (B) and astrocytes outside the BgL (C) upon ischemic injury in *Nes-Td* mice (B: Student’s t-test, p<0.02, n≥3). D) Experiential timeline for *Hopx-Td* GIFM. **E-G)** Quantification of the density of TdTom+ Bg (F) and astrocytes outside the BgL (G) shows a significant increase in the density of TdTom+ Bg upon injury (Student’s t-test, p<0.0118, n=4) but no change in astrocytes (Student’s t-test, p>0.05, n=4). **H)** The majority of the TdTom+ Bg were S100β negative (Two-way ANOVA, F_(1,4)_=222, p<0.0001, n=3). **I)** Summary of findings and the working model for the relationship between Bg, *Hopx*+ Bg and *Nes+* Bg. The curved lines indicate proliferation, the solid lines show confirmed relations between different Bg types, and the thickness of the lines indicates the increase upon injury and/or SAG with respect to homeostasis. The grey lines indicate the trends that were observed but not significant. The dashed line represents indirect evidence, and the black dotted lines show additional potential mechanisms that were not addressed in this study. Multiple comparison tests are shown in the figures. Scale bars: 200 µm

## Discussion

Here, we characterised a rare subpopulation of Bg that expresses *Nes*. These cells largely do not express mature astrocyte markers and have a higher in *vitro* neurosphere formation ability compared to *Nes*-cells, highlighting a stem/progenitor cell-like nature. Importantly, their density significantly increased with injury or SHH activation, and the two stimuli act together to further increase the density of Nes-CFP+ Bg following two cerebellar injuries. However, only a small proportion of the Nes-CFP+ cells are proliferative (10% incorporate EdU during a 1 hr pulse), and this proportion does not change with injury or SAG activation. This suggests that some *Nes-* Bg, perhaps mainly those expressing *Hopx*, become *Nes+* upon SAG administration and/or injury to the adult cerebellum. Our bulk ATAC-seq comparison of adult and neonatal Nes-CFP+ cells revealed the silencing of stem cell proliferation and differentiation programmes during development, which likely contributes to the reduced regenerative potential of adult Bg. Moreover, GIFM analysis shows that *Nes+* Bg do not generate new neurons after injury to the adult cerebellum. Finally, our results highlight the cellular heterogeneity of Bg in the adult cerebellum with respect to transcriptional identity and their response to injury or SHH signalling. Further research is needed to unravel the molecular drivers of these cellular differences and determine if there are ways to stimulate neuron production.

Analysis of the distribution of the Nes-CFP+ cells across the cerebellum revealed some interesting observations. These cells reside across the medial-lateral axis of the adult cerebellum, and the majority are located in the BgL (*Nes+* Bg). Interestingly, we observed a significantly higher density of *Nes+* Bg in the posterior than the anterior vermis. While the relevance is unclear, one possibility is that the posterior *Nes+* Bg receive additional secreted signals from the CSF and 4^th^ ventricle choroid plexus^41^, possibly including more SHH, that promote *Nes* expression in a higher proportion of Bg. The posterior sector was not analysed in the injury analysis as the ischemic injury did not target this region; therefore, how the posterior *Nes+* Bg cells respond to injury remains unknown.

Our GIFM shows that the *Nes+* Bg remain as glia following injury. A question remains as to whether *Nes+* Bg represent a subgroup of neonatal gliogenic NEPs that had expressed *Hopx* but did not undergo full differentiation to become S100β+. Previous studies suggest that adult neural stem cells in the forebrain neurogenic niches are set aside during development^42,43^. When the *Nes+* Bg are born remains to be determined. Importantly, whether *Nes* expression in the BgL marks a specific stable subtype or a dynamic cell state of Bg under different physiological conditions remains unknown.

Although the neonatal cerebellum is highly regenerative and the *Hopx*-expressing gliogenic-NEPs can undergo adaptive reprogramming upon injury at birth to become GCPs, the adult cerebellum is not spared from the overall stunted regenerative capacity of the rest of the adult brain^44^. Our chromatin accessibility data demonstrate that Nes-CFP+ Bg in the adult cerebellum do not turn on regenerative programmes upon injury. Perhaps such mechanisms are in place to prevent aberrant reprogramming, which could lead to tumorigenesis.

SHH is a crucial mitogen for cerebellum development. Once postnatal development ends, Purkinje cells continue to secrete SHH, however, its function in the adult cerebellum remains poorly understood. Our data indicate that at steady state, most adult Bg are *Gli1*+, suggesting some level of SHH pathway activity. However, only a subset of Bg become *Nes+* upon increased SHH activity. Why only some Bg are able to become *Nes*+ upon SHH activation (and/or injury) remains to be determined. Importantly, during neonatal cerebellum development, SHH promotes the proliferation of NEPs and GCPs; however, upon an increase in SHH signalling in the adult cerebellum, the percentage of Nes-CFP+ Bg that proliferate (incorporate EdU) do not increase, although overall, the number of Nes-CFP+ Bg increases, and thus there are more proliferating cells. In contrast to the cerebellum, analysis of SHH activity in the adult forebrain showed that only the quiescent neural stem cells respond to SHH (express *Gli1*) in the neurogenic niches^45^. Furthermore, SHH signalling is required for the maintenance of the quiescent neural stem cells in the postnatal brain^46,47^ and deletion of *Patched* in the adult subventricular zone NSCs, which upregulates HH signalling, leads to a depletion of the proliferating NSCs and accumulation of quiescent cells^48^. This raises the possibility that SHH signalling functions differently in the adult Bg compared to its mitogenic role during postnatal cerebellar development.

The lack of neuron production upon injury and SAG administration reveals that more extensive epigenetic changes need to be reversed in adult Bg compared to in neonates. In addition to the cell-intrinsic programmes, changes in the brain microenvironment also likely influence the regenerative potential. For example, microglia have been shown to inhibit the direct reprogramming of the Müller glia in the retina into neurons^49^, whereas our analysis in the neonatal cerebellum suggests that microglia are proregenerative^50^. Given that our injury models also show an increase in microglia activation and GFAP levels (Figure 2B-D and Supplementary Figure 1C), indicating reactive gliosis, how these cells influence *Nes*+ Bg behaviours should be studied.

Collectively, our results shed light on the injury response of the adult cerebellum, with a focus on the *Nes+* Bg during homeostasis and upon injury. Importantly, our results provide insights into the reasons behind the declining regenerative potential of the orthogonal *Nes*-expressing populations that arise from the neonatal cerebellum. Finally, activation of developmental pathways, specifically SHH, can promote an enhanced response to injury via increasing the number of *Nes*+ Bg. However, SHH activation is not sufficient to induce neuron production, suggesting a requirement for additional neurogenic cues. Our results highlight the importance of understanding the developmental signalling that stimulates adaptive reprogramming in the neonatal cerebellum as a means to understand why the adult cerebellum cannot regenerate, and to identify ways to stimulate regeneration safely in the adult brain.

## Materials and Methods

### Animals

All the mouse experiments were performed according to protocols approved by the Memorial Sloan Kettering Cancer Center Institutional Animal Care and Use Committee (IACUC, protocol no 07-01-001). Animals were housed on a 12-hour light/dark cycle and given access to food and water *ad libitum*.

The following mouse lines were used: *Nes-Cfp*^4,51^, *Hopx^CreERT2^* ^52^, *Nes-FlpoERT2^4^ (*Stock no: 040094, The Jackson Laboratories), *Rosa26^FRT-STOP-FRT-TdTomato^*(Stock no: 021875, The Jackson Laboratories)^53^*, Rosa26^lox-STOP-loxTdTomato^* (ai14, Stock no: 007909, The Jackson Laboratories)^54^, *Gli1^eGfp^* (Stock no: 040090, The Jackson Laboratories)^38^, *P1-Shh*^22,55^. Animals were maintained on an outbred Swiss Webster background. Both sexes were used for analyses (except for bulk ATAC-seq, explained below), and experimenters were blinded for genotypes whenever possible.

Tamoxifen (200µg/g, Sigma) was injected intraperitoneally into 4-6 week old mice once or thrice every other day. EdU (50µg/g, Life Technologies). The SMO agonist (SAG,100µg/g, Tocris) was injected intraperitoneally at the described timepoints (Figure 4A, C). Diphtheria Toxin (30ng/g, List Biological Laboratories Inc.) was injected subcutaneously or intraperitoneally into the neonatal or adult *GC-DTR* mice, respectively. Pregnant *GC-DTR* dams were given Doxycyline (0.02 mg/mL in drinking water, Sigma) between E8.5-12.5 to induce GCP-specific recombination as previously described^23^.

### Photothrombotic Injury

Ischemic stroke was performed similar to a previously described protocol with minor modifications and using aseptic technique ^34^. Briefly, in order to induce local stroke, 4-6 week old mice were anaesthetised (Ketamine (100mg/kg)/Xylaxine (10mg/kg)). Following intraperitoneal administration of 100µL of 20mg/mL Rose Bengal (Sigma), animals’ heads were shaved. A small incision over the cerebellum was made, and a thin cold LED (Leica) light beam (2-3mm in diameter) was shone over the vermis area for 15 minutes. The incision was sutured, and the animals were taken to recovery. Animals were provided with necessary analgesics (0.1mg/kg Buprenorphine) during recovery.

### Tissue Preparation and Histological Analysis

Animals (4 weeks or older) were perfused with ice-cold PBS followed by 4% PFA, following anaesthesia. After dissection, brains were fixed further for an additional 24-48 hours in 4% PFA in PBS. Fixed brains were switched to 30% Sucrose in PBS until sinking. Then, the brains were embedded in OCT (Tissue-Tek) for cryosectioning. 14um-thick sections were obtained using a cryostat (Leica, CM3050S) and stored at -20°C.

Immunofluorescent analysis was performed on cryosections. Slides were allowed to warm to room temperature (RT) and washed once with PBS. 1 hour blocking was performed at RT using 5% Bovine Serum Albumin (BSA, Sigma) in PBS-T (PBS with 0.1% Triton-X). Slides were incubated with primary antibodies diluted in blocking solution at 4°C overnight (Supplementary Table 3). Slides were then washed with PBS-T (3 x 5 min) and incubated with fluorophore-conjugated secondary antibodies (1:500 in blocking buffer, Invitrogen). Hoechst 33258 (Invitrogen) was used to label the nuclei and the slides were mounted with Fluoro-Gel mounting media (Electron Microscopy Sciences).

To detect EdU, a Click-it EdU assay with Sulfo-Cyanine5 azide (Lumiprobe corporation, A3330) was used.

### Imaging and Image Analysis

Images were collected either with a DM6000 Leica microscope, Zeiss LSM 880 confocal microscope or Nanozoomer (Hamamatsu) and processed using ImageJ Software (NIH). For each quantification, three midline parasagittal sections/brain were analysed and data were averaged, except for Nes-TD brains, where five sections/brain were analysed and the data were summed due to the low frequency of labelling. Cells were counted using the Cell Counter plugin for ImageJ (NIH). Categorization of cells as Bg or astrocytes was based on morphology. For the ischemic injury model, the vermal central zone was quantified and for other quantification, lobule 4/5 was assessed, except for the pan cerebellar analysis presented in Figure 1, where all lobules at all medial and lateral levels were quantified.

### Cell dissociation for FACS

*Nes-Cfp/+* cerebella were dissected into ice-cold 1x Hank’s Buffered Salt Solution (Gibco) and dissociated using Accutase (Innovative Cell Technologies) at 37°C for 10-15 min. After dissociation, Accutase was diluted with 3x excess volume of neural stem cell media (Neurobasal medium, supplemented with N2, B27 (without vitamin A)), and nonessential amino acids (Life Technologies, Gibco) and cells were filtered using a 40 µm mesh cell strainer. Following filtering through a cell strainer and trituration in media to single cells, cells were layered over a 5mL density gradient (albumin-ovomucoid inhibitor solution, Worthington) and centrifuged at 70g for 6 minutes to remove debris, followed by an additional 5 min of centrifugation in a chilled centrifuge at 500g twice. The final pellet was resuspended in neural stem cell media and strained using strainer cap tubes (Falcon) for downstream experimentation. All centrifugation was performed at 4°C and cells were kept on ice when possible. A BD FACS Aria II (BD Biosciences) was used for cell isolation, and cells were sorted using the 100 μm nozzle for other downstream analyses.

### Limiting dilution neurosphere formation assay

FACS-isolated Nes-CFP+ and Nes-CFP-cells were plated at three densities (60, 600 and 6,000 cells/well) in Ultra-low attachment 96-well plates (Corning) using Neurobasal media (Gibco, Life Technologies), supplemented with N2 (Gibco, Life Technologies), B27 (without vitamin A) (Gibco, Life Technologies), L-Glutamine (Gibco, Life Technologies), non-essential amino acids (Gibco, Life Technologies) and 20 ng/mL of recombinant EGF (Gibco, Life Technologies) and 20 ng/ml FGF2 (Gibco, Life Technologies). Growth factors were supplemented every other day, and the number of neurospheres was analysed 7 days after plating. For serial passaging, Neurospheres were dissociated every 7 days using Accutase (Innovative Cell Technologies) and cells were replated at the original density (600 cells/well).

### Bulk ATAC-seq

#### Sample preparation

FACS-isolated Nes-CFP+ cells (30,000-50,000 per replicate) were isolated from male neonatal control or P1 DT-injected P5 *GC-DTR* mice and adult P30 control or DT-injected adult mice that were collected 4 days after injury. 2-3 cerebella were pooled for each neonatal sample, and 1-2 cerebella were pooled for each adult CB. 2 replicates were prepared for each condition. Cells were immediately processed for nuclei preparation and transposition using the OMNI-ATAC protocol ^56^. Sequencing was performed at the MSKCC genomics core facility using the Illumina NovaSeq S4 platform.

Raw sequencing reads were 3’ trimmed and filtered for quality and adapter content using version 0.4.5 of TrimGalore (https://www.bioinformatics.babraham.ac.uk/projects/trim_galore), with a quality setting of 15, and running version 1.15 of cutadapt and version 0.11.5 of FastQC. Version 2.3.4.1 of bowtie2 (http://bowtie-bio.sourceforge.net/bowtie2/index.shtml) was used to align reads to mouse assembly mm10 and alignments were deduplicated using MarkDuplicates in version 2.16.0 of Picard Tools. Enriched regions were discovered using MACS2 (https://github.com/taoliu/MACS) with a p-value setting of 0.001, filtered for blacklisted regions (http://mitra.stanford.edu/kundaje/akundaje/release/blacklists/mm10-mouse/mm10.blacklist.bed.gz), and a peak atlas was created using +/- 250 bp around peak summits. The BEDTools suite (http://bedtools.readthedocs.io) was used to create normalized bigwig files. Version 1.6.1 of featureCounts (http://subread.sourceforge.net) was used to build a raw counts matrix and DESeq2 was used to calculate differential enrichment for all pairwise contrasts. Peak-gene associations were created by assigning all intragenic peaks to that gene, while intergenic peaks were assigned using linear genomic distance to the transcription start site. Network analysis was performed using the assigned genes to differential peaks by running enrichplot::cnetplot in R with default parameters. Motif signatures were obtained using Homer v4.5 (http://homer.ucsd.edu) on differentially enriched peak *regions*.

### Statistical comparisons

Prism (GraphPad) was used for all statistical analysis. Statistical comparisons used in this study were Student’s two-tailed t-test; One-way and Two-way analysis of variance (ANOVA), followed by post hoc analysis with Tukey’s or Sidak’s test for multiple comparisons. Sample size, relevant F-statistics and p-values are stated in the figure legends. The p-values of the relevant post hoc multiple comparisons are shown in the figures. All t-tests were two-tailed.

The statistical significance cutoff was set at p<0.05. Data is presented as mean ± standard deviation (SD) of the mean unless otherwise stated in the figure legend.

## Data availability

Bulk ATAC-seq data has been submitted to GEO (Accession ID: GSE306093).

## Acknowledgements

We thank the past and present members of the Joyner lab for discussions, the Bayin lab members for reviewing the manuscript, and the MSKCC Genomics Facility and Epigenomics Core and the MSKCC Flow Cytometry core facility for technical services and support.

## Author Contributions

NSB: conception, design, data acquisition, analysis, and interpretation, writing of the manuscript and funding acquisition; DNS: data acquisition; RK: epigenomic analysis; ALJ: conception and design of the project, data interpretation, writing of the manuscript and funding acquisition. All authors reviewed the manuscript.

