## Supplementary Figures and Legends for "The regenerative potential of adult *Nestin+* cerebellar astroglia is limited compared to in neonates"

**Supplementary figure 1**

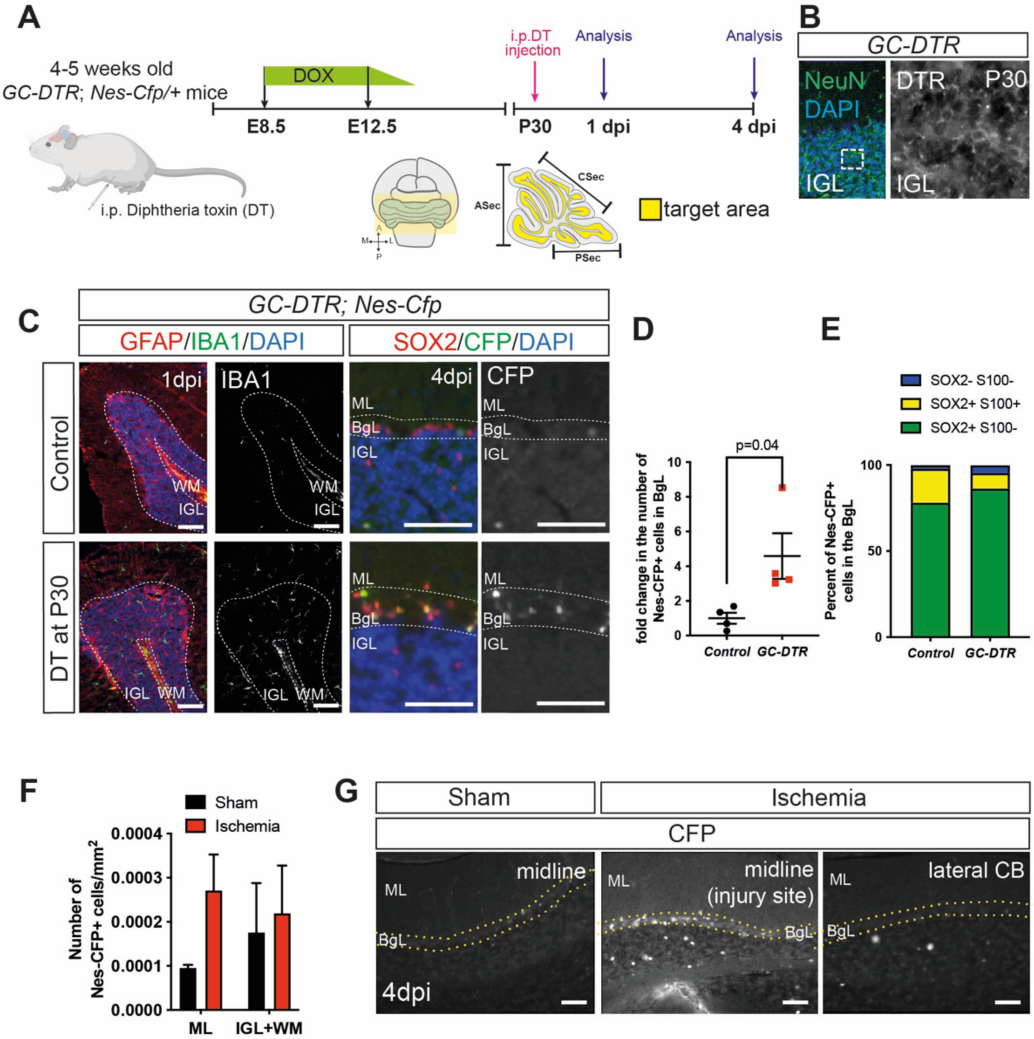


**Supplementary Figure 1. Nes-CFP+ Bg density is increased in response to granule cell ablation in the adult cerebellum, and the effects of ischemia are restricted to the injury site. A)** Schematics explaining the pharmacogenetic granule cell ablation model (*GC-DTR*). **B)** Immunofluorescent analysis of IGL shows DTR expression in the granule cells in *GC-DTR* animals. **C)** Immunofluorescent analysis of GFAP+ and IBA+ cells upon injury and the increase in Nes-CFP+ Bg 4 days after ablation. **D-E)** Quantification of the Nes-CFP+ Bg shows a significant increase in the density of cells upon injury (Student's t-test, p=0.04, n=4) and suggests that the increase is driven by the S100β- population (n=4). **F)** Density of the Nes-CFP+ cells in the other layers of the cerebellar cortex shows high variability upon injury and no significant changes (ischemic model). **G)** Immunofluorescent analysis of the injury site and lateral cerebellum (away from the injury) shows that the increase in the Nes-CFP+ Bg density is specific to ischemic injury. Scale bars: 100 μm

### Supplementary Figure 2

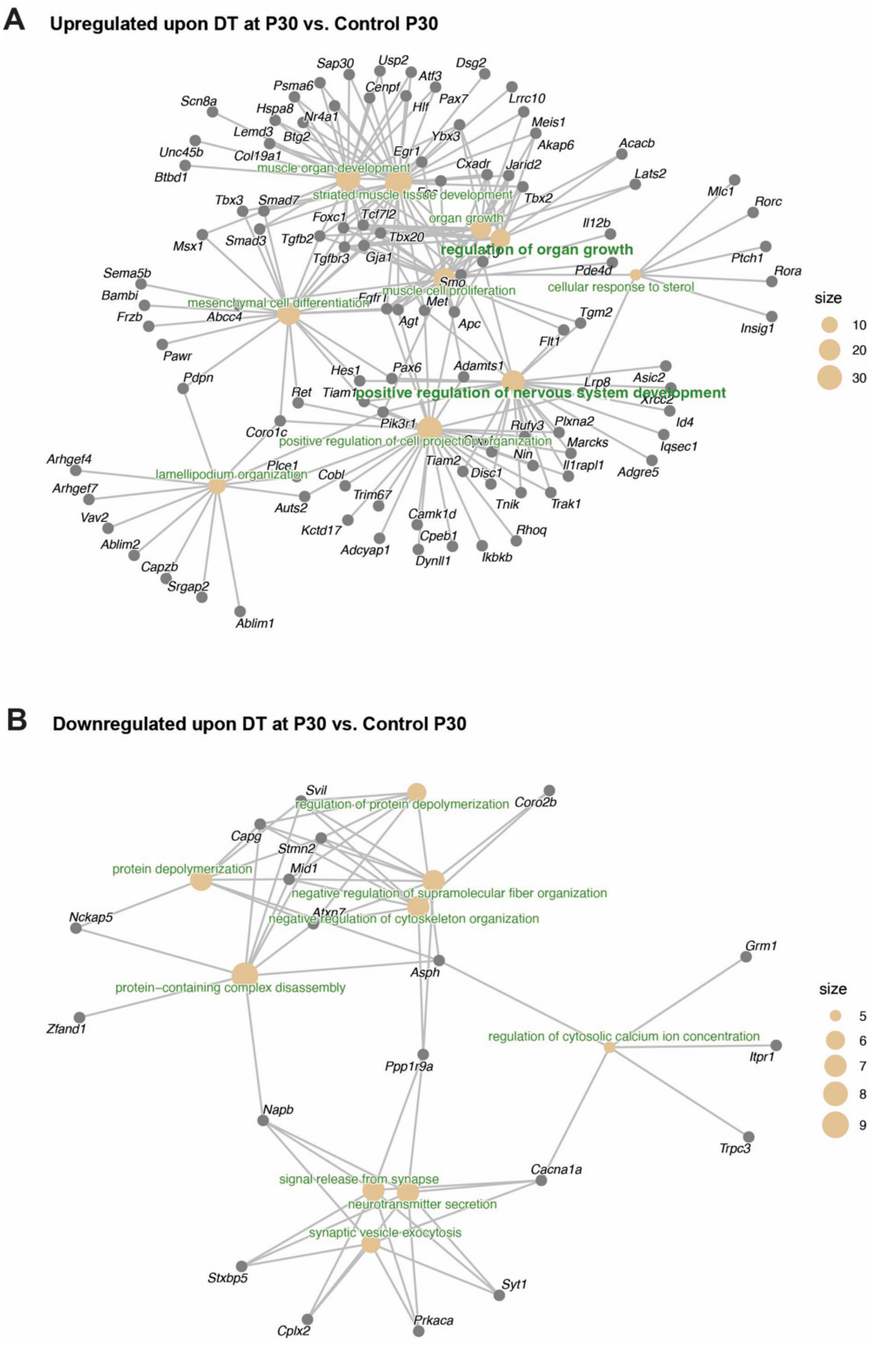


**Supplementary Figure 2. Injury to the adult cerebellum causes significant changes to the chromatin landscape of Nes-CFP+ cells in *GC-DTR* mice compared to the controls. A-B)** Network analysis of genes that are associated with top differentially open chromatin in response to injury at P30 (A, n=575 genes) or closed (B, n=160 genes) and the associated gene ontology terms associated with these genes (fold change=1.5, FDR=0.2, see Supplementary Table 1).

### Supplementary Figure 3

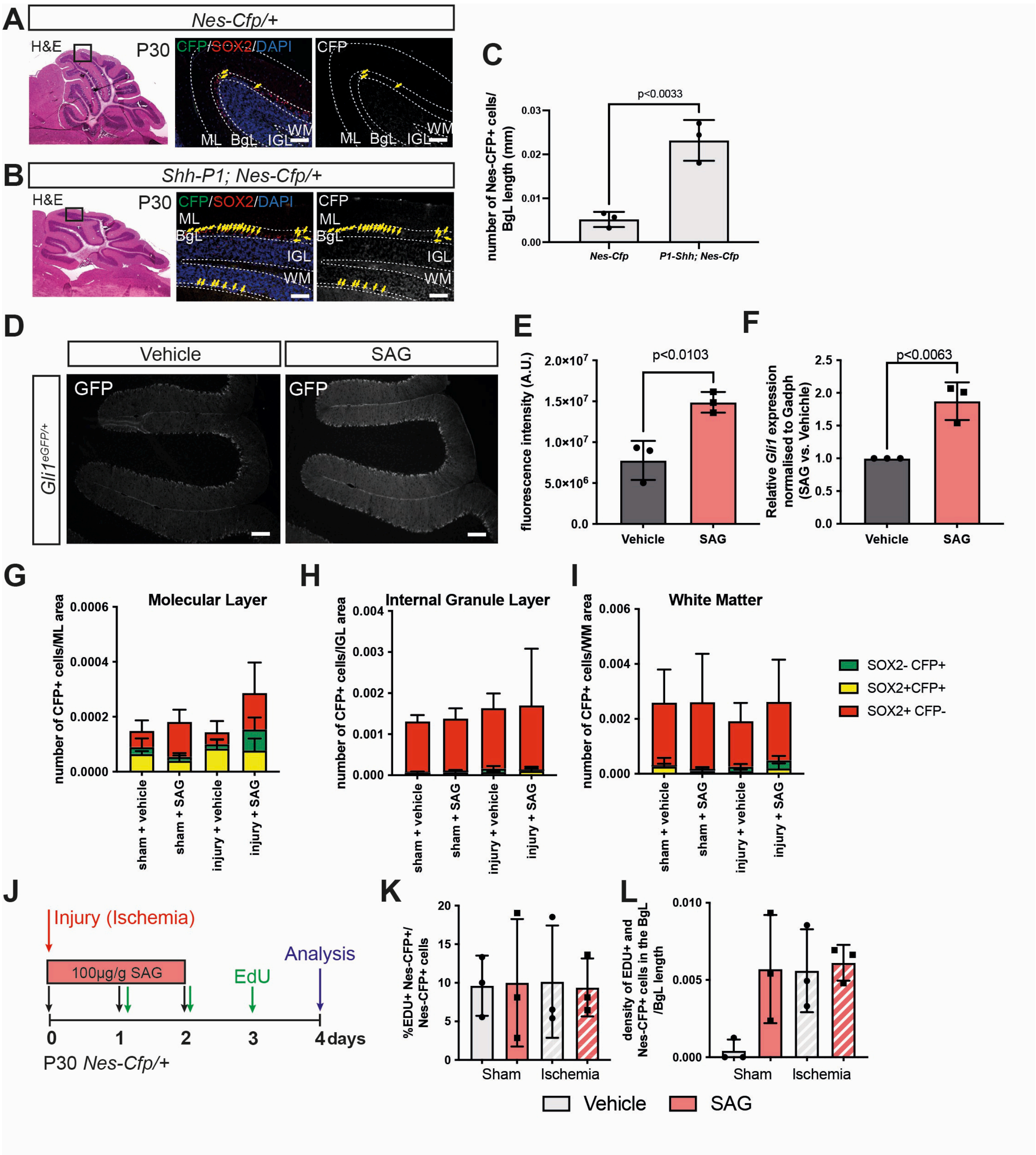


**Supplementary Figure 3. Activation of SHH signalling using a transgene or small molecule agonist against SMO. A-C)** Analysis of *Shh-P1; Nes-Cfp* transgenic animals shows that an extra 3rd copy of the mouse *Shh* gene promotes increased density of the Nes-CFP+ Bg (Student's t-test, p<0.0033, n=3). **D-F)** Activation of SHH signalling using SAG shows significantly increased GFP intensity in the BgL and ML of a *Gli1*$^{eGFP/+}$ reporter mouse line (Student's t-test, p<0.0103, n=3) and a significant upregulation of *Gli1* transcripts 6 hours after a dose of 10μg/g SAG injection (Student's t-test, $p<0.0063$, $n=3$). **G-I)** Quantification of Nes-CFP+ and SOX2+ cells outside the BgL shows no significant changes upon SAG and/or ischemic injury, highlighting that the Bg are the primary responding population to both stimuli. **J.** Experimental timeline. **K-L)** Quantification of the percentages (K) and density (L) of EdU+ Nes-CFP+ Bg upon SAG and/or ischemic injury (Two-way ANOVA, $p>0.05$, $n=3$). Scale bars: 100 μm, except for D: 200 μm

### Supplementary Figure 4

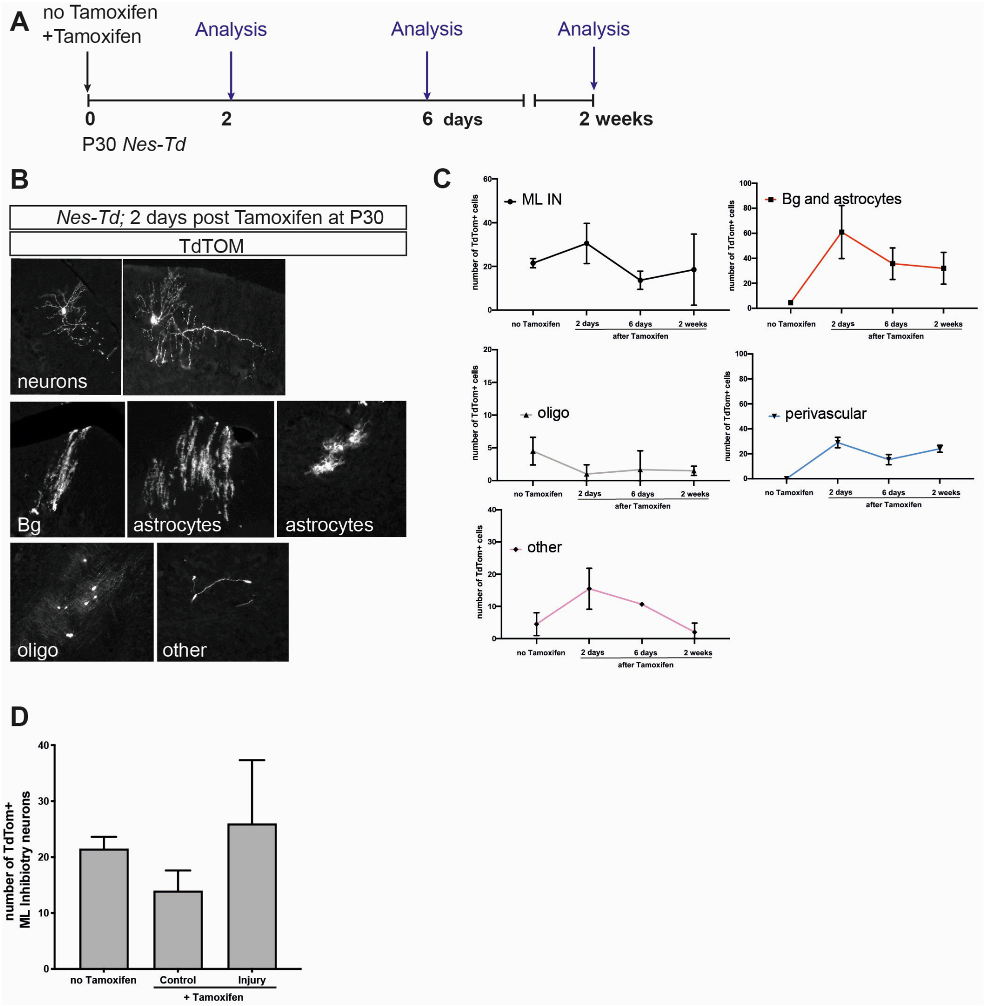


**Supplementary Figure 4. Analysis of *Nes-Td* brains with or without Tamoxifen injection over time highlights the non-specific labelling of some cerebellar cell types. A)** Experimental plan. **B)** Examples of some TdTom labelled cell types in the adult cerebellum and how the different cell shapes were used for annotation. **C)** Quantification of the number of TdTom+ cells categorised by the cell types. Note that other than astroglia (Bg and astrocytes) and perivascular cells, all other populations showed equal amounts of labelling without and with tamoxifen. **D)** Injury does not lead to significant changes in the density of TdTom+ ML inhibitory neurons two weeks after injury (Tamoxifen administered 4 days after injury) compared to the brains that were not injected with Tamoxifen, or the brains that were injected with Tamoxifen but were not subjected to injury.

**List of supplementary tables**

**Supplementary Table 1**. Differentially open chromatin in Nes-CFP+ cells between control and injured adult *GC-DTR* brains (for significant results, a threshold of fold change=1.5, FDR=0.2 was used).

**Supplementary Table 2.** Differentially open chromatin between P5 and P30 Nes-CFP cells (only from controls) (for significant results, a threshold of fold change=2, FDR=0.1 was used).

**Supplementary Table 3**. List of antibodies

| Target | Catalog Number | Company | Dilution |
|---|---|---|---|
| goat α-SOX2 | AF2018 | R&D System | 1/200 (adult)-<br>1/500 (pups) |
| rat α-GFP (CFP) | 04404-84 | Nacalai Tesque | 1/1000 |
| rabbit α-S100B | Z0311 | DAKO | 1/1000 |
| mouse α-NeuN | MAB3777 | Millipore | 1/1000 |
| rabbit α-IBA1 | HPA030180 | Sigma (Prestige Ab) | 1/1000 |
| goat α-hHB-EGF(DTR) | AF231 | R&D System | 1/500 |
| chicken α-GFAP | Ab4674 | Abcam | 1/500 |
| rabbit α-IBA1 | 019-19741 | Wako Chemicals | 1/500 |
